# Functional divergence of conserved developmental plasticity genes between two distantly related nematodes

**DOI:** 10.1101/2024.08.29.610235

**Authors:** Sara Wighard, Hanh Witte, Ralf J. Sommer

**Affiliations:** Department for Integrative Evolutionary Biology, Max Planck Institute for Biology, Max-Planck-Ring 9, 72076 Tübingen, Germany; Institute of Molecular Biotechnology of the Austrian Academy of Sciences (IMBA), Vienna BioCenter (VBC), 1030 Vienna, Austria

**Keywords:** Polyphenisms, developmental plasticity, *Allodiplogaster sudhausi*, *Pristionchus pacificus*, eud-1/sulfatase, nuclear-hormone-receptors, gene divergence

## Abstract

Genes diverge in form and function in multiple ways over time; they can be conserved, acquire new roles, or eventually be lost. However, the way genes diverge at the functional level is little understood, particularly in plastic systems. We investigated this process using two distantly related nematode species, *Allodiplogaster sudhausi* and *Pristionchus pacificus*. Both these nematodes display environmentally-influenced developmental plasticity of mouth-form feeding structures. This phenotype can be manipulated by growth on particular diets, making them ideal traits to investigate functional divergence of developmental plasticity genes between organisms. Using CRISPR-engineered mutations in *A. sudhausi* mouth-form genes, we demonstrate examples of the various ways ancestral genes regulate developmental plasticity and how these roles can progressively diverge. We examined four ancestral genes, revealing distinct differences in their conservation and functional divergence in regulating the mouth phenotype in both species. Specifically, certain genes retain the same characteristics, while others have acquired a new function. Additionally, two ancestral genes retain their functions as switch genes, which completely prevent a phenotype, and the other two display quantitative effects, with knockouts in these genes displaying intermediate phenotypes. Remarkably, despite the evolutionary distance, all genes examined were involved in mouth-form regulation. Finally, multiple gene knock-out mutants were engineered, with key sulfatase-encoding genes acting downstream of all others, suggesting they play a major role in mouth-form plasticity. Together, this study represents the first mutant-based functional analysis of the evolution of developmental plasticity between two highly diverged species, offering new insights into the genetic mechanisms underlying phenotypic evolution.

**Article Summary:** While evolutionary divergence of genes is well-studied at the sequence level, the resulting functional and phenotypic consequences are less known, particularly in plastic systems. Here, we examined functional divergence of a set of genes involved in developmental plasticity of mouth-form between two highly diverged nematode species. We found that all studied genes control mouth-form plasticity in both species; however, with strong functional divergence and gene-specific quantitative effects or even novel functions. Thus, there is a spectrum from full conservation, partial conservation to the gain of a new function; with genes involved in sulfation showing the strongest conservation during evolution.

## Introduction

Over time, genes diverge from their ancestral version in both sequence and function. Numerous studies have estimated the timing of divergence, often using substitution rates along with fossil records (Pinho and Hey 2010; Tamura *et al*. 2012; Smith *et al*. 2018). These are continuously being updated and optimised, enabling us to better understand the gradual changes in DNA and amino acid sequences. However, it is also vital to examine changes at the phenotypic (functional) level. Here, we compare two distantly related nematodes, *Pristionchus pacificus* and *Allodiplogaster sudhausi*, who nonetheless belong to the same superfamily, the Diplogastridae (Fig. 1a) (Sudhaus and Fürst von Lieven 2003). Both these species display developmental (phenotypic) plasticity, the phenomenon whereby organisms with the same genotype can form different phenotypes based on environmental cues. Recent work in the likes of spadefoot toads (Pfennig *et al*. 2010), horned beetles (Moczek 1998) and indeed diplogastrid nematodes (Sommer 2020) has shown developmental plasticity plays a key yet under-appreciated role as a driver of evolutionary events and novelty.

**Fig. 1.**
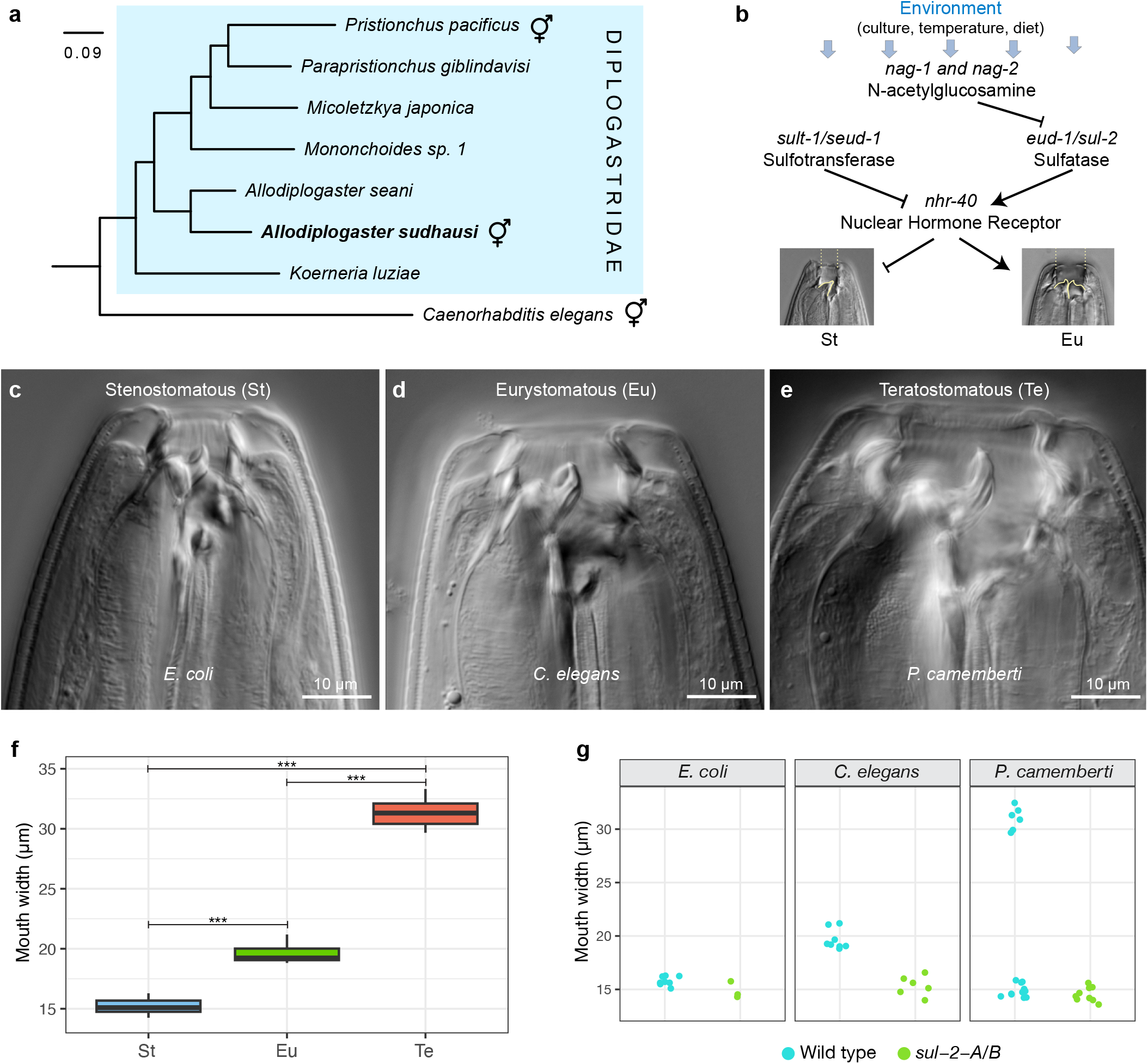
The nematode *Allodiplogaster sudhausi* can form three potential adult morphs, with conserved sulfatase genes regulating mouth-form phenotype. a) *A. sudhausi* belongs to the Diplogastridae nematode family and is a distant relative of *Pristionchus pacficus*. Hermaphroditic symbols indicate species that are androdeiocious (hermaphrodite/male). b) The gene regulatory network controlling adult mouth-form phenotype has been well-elucidated in *P. pacificus*, with many genes identified that act as a switch in becoming either Stenostomatous (St) or Eurystomatous (Eu). c - e) *A. sudhausi* adults can form three possible mouth-form morphs, as indicated by differential interference contrast (DIC) microscopy pictures of (c) ‘narrow-mouthed’ St from an *E. coli* diet, (d) ‘wide-mouthed Eu’ from predation on *C. elegans* and (e) ‘monster-mouthed’ Teratostomatous (Te) from a fungal *P. camemberti* diet. f) Mouth width measurements show there are significant differences between St, Eu and Te (Kruskal-Wallis, ***P < 0.001; pairwise Wilcox test, P < 0.01 for each comparison; n = 8 to 12 biological replicates). g) Double mutant knock-outs in the sulfatases, *sul-2-A/B* remain St under all three diets, in stark contrast to wild type worms. Data and figures obtained from previous work (Wighard et al. 2024).

Members of the Diplogastridae display developmental plasticity in mouth-form phenotype, with most species capable of forming two alternative adult mouth-form morphs that are adapted to different environmental conditions (Susoy et al. 2015a). For example, *P. pacificus* can form the narrow-mouthed stenostomatous (St) morph which lacks a sub-ventral tooth and is bacterial-feeding, or it can form the wide-mouthed eurystomatous (Eu) morph which has two teeth and can predate on other nematodes (Bento *et al*. 2010). The mechanisms behind this plasticity have been well-studied, with important genes identified that regulate the two mouth-form phenotypes (Ragsdale *et al*. 2013; Sieriebriennikov *et al*. 2018; Namdeo *et al*. 2018; Bui *et al*. 2018; Levis and Ragsdale 2023) (Fig. 1b). We recently examined *A. sudhausi*, which diverged early in the Diplogastridae lineage and displays mouth-form plasticity. While *A. sudhausi* still retains the equivalent St and Eu morphs that are found in most diplogastrids, we were surprised to discover it can form three possible morphs (Fig. 1c - f). The newly identified third morph, termed teratostomatous (Te), has the widest mouth and appears when grown on the fungus *Penicillium camemberti* (Wighard et al. 2024). Interestingly, this Te morph, in contrast to St or Eu individuals, exhibits cannibalism against genetically identical kin, a new behaviour in nematodes (Wighard et al. 2024). Taken together, both the *P. pacificus* and *A. sudhausi* mouth-forms occur as a polyphenism, *i*.*e*. produce discrete alternative phenotypes. This polyphenic trait enables us to compare discrete characteristics between species and is ideal to examine the functional changes of ancestral genes over time.

Multiple mouth-form genes have already been identified in *P. pacificus*, based on previous studies that used forward and reverse genetics to generate knock-out mutants (Fig. 1b). For instance, the sulfatase-encoding gene *eud-1* acts as a developmental switch (Ragsdale *et al*. 2013) and is part of a supergene locus that also contains two α-N-acetylglucosaminidase (*nag*) genes. While *eud-1* mutants can only form the St morph, *nag-1 nag-2* double mutants have the opposite phenotype and are constitutively Eu (Sieriebriennikov *et al*. 2018). This supergene locus recently evolved, is subject to natural variation (Dardiry *et al*. 2023) and acts as a major regulator of mouth-form plasticity. Additionally, the nuclear hormone receptor *nhr-40*, considered a target of *eud-1*, plays an important role in mouth-form regulation (Sieriebriennikov *et al*. 2020), as does the sulfotransferase-encoding gene *sult-1* (also known as *seud-1*) (Namdeo *et al*. 2018; Bui *et al*. 2018). In *P. pacificus*, mutations in these genes result either in an all-St phenotype (*eud-1; nhr-40*), or an all-Eu phenotype (*nag-1 nag-2; sult-1*). Therefore, these genes are ideal candidates to examine the divergence of homologs between both *P. pacificus* and *A. sudhausi*. Importantly, functional studies of homologs of *eud-1* were previously performed in *A. sudhausi* (Wighard et al. 2024). There, we found two duplicate genes, termed *sul-2-A* and *sul-2-B*. These sulfatase gene(s) act as a switch in both species and regulate the Eu morph, with knock-outs remaining St across all conditions (Fig. 1g). Notably, in *A. sudhausi* the two genes additionally regulate the third Te morph, indicating the additional morph may have evolved by co-option of the existing Eu machinery (Wighard et al. 2024). Thus, the sulfatase genes display conservation of function between both species in Eu regulation; however, they also evolved a novel function in Te regulation in *A. sudhausi*. We therefore wanted to examine other mouth-form genes to see if they show similar conservation between both species, or whether there is further novelty to be found.

Estimating the divergence between *P. pacificus* and *A. sudhausi* is crucial to determine the time that has passed, which allows evolutionary changes to take place and mouth-form genes to diverge. Amber fossil records of diplogastrids as well as other nematodes have been found (Poinar 2011), with the oldest known diplogastrid taxon estimated to be from 99 million years ago (mya). Phylogenomic analysis taking the fossil dating and divergence estimates in account have recently been evaluated by Qing *et al*. (2023). They performed divergence time dating on different nematode clades using molecular dating in comparison to the Tardigrada sister group. The *P. pacificus* and *A. sudhausi* genomes were both included in these analyses, with *A. sudhausi* estimated to have diverged from their common ancestor approximately 180 mya in the Jurassic period. This is an exceedingly long time for evolutionary changes to happen, particularly when one takes into account the short generation times of nematodes; four days for *P. pacificus* (Schroeder 2021) and eight days for *A. sudhausi* (Fürst von Lieven 2008). Evaluating plasticity in both these species should give an indication of the level of selection on mouth-form genes and allow us to see if they retain the same function.

*A. sudhausi* and *P. pacificus* are suitable comparative species for examining conservation of gene function due to the following traits: 1) self-fertilising hermaphroditism, 2) the availability of assembled and annotated genomes, 3) a large evolutionary distance between both species, 4) the retention of mouth-form plasticity and 5) the availability of genetic knock-out tools (Wighard et al. 2022). Importantly, while mouth-form plasticity was conserved between both species, novelty arose in *A. sudhausi* with the formation of the Te morph. This enables us to examine how evolution in plasticity had an impact on, or was impacted by the underlying genes. Importantly, we can manipulate the mouth-form phenotype by using three different diets (Wighard et al. 2024). Specifically, *A. sudhausi* becomes St on *Escherichia coli* bacteria, Eu on *C. elegans* nematodes and Te on *P. camemberti* fungus. The phenotype is consistent for the first two diets, while on *P. camemberti*, worms become either St or Te, with density increasing Te occurrence. These different diets and their respective mouth-form phenotypes enable us to reproducibly examine the mutant phenotypes on different conditions.

The goal of this project was to knock-out putative mouth-form genes in *A. sudhausi* and determine whether their role in mouth-form plasticity and the manner of gene regulation is conserved in both *A. sudhausi* and *P. pacificus*. The presence of the third mouth-form in *A. sudhausi* additionally enabled us to examine the role these genes may play in a novel trait. It is important to note that *A. sudhausi* recently underwent whole genome duplication (WGD) (Wighard et al. 2022), meaning that most genes have duplicate copies. However, due to the recency of the event, the majority of genes are redundant as there has been too little time for divergence between the copies. Therefore, two *A. sudhausi* genes needed to be targeted for every one homologous gene in *P. pacificus*.

Here, we targeted six additional genes in *A. sudhausi*, and created combinations of mutant knock-out lines in order to compare the ways in which mouth-form genes showed conservation or divergence between *A. sudhausi* and *P. pacificus*. We found that knock-outs in all *A. sudhausi* homologous mouth-form genes produced mutants with mouth phenotypes that differed from wild type and are thus involved in mouth-form regulation. However, the means in which genes controlled plastic traits often differed between species, with homologous genes displaying unique profiles, indicating distinct roles in the divergence of these genes from their ancestral forms.

## Results and Discussion

A single guide RNA (gRNA) targeting each pair of WGD-derived genes was injected into wild type *A. sudhausi* worms to generate knock-out mutants (Fig. 2a). Homozygous frameshift mutations in the genes of interest were then selected for in the subsequent progeny. Both single and double mutant knock-outs were generated, although single mutants continuously displayed redundancy and a wild type phenotype, meaning double mutants were instead analyzed in-depth. These mutant lines were then grown up from eggs via a bleaching protocol (Hope 1999) on the three different diets. In the wild type line, these diets have consistent and replicable phenotypes on: 1) *E. coli*, which always becomes St, 2) *C. elegans*, which always becomes Eu and 3) *P. camemberti*, which becomes St or Te, depending on population density (Wighard et al. 2024). This manipulation of diet allowed us to investigate the influence of mutant knock-outs on the St, Eu and Te phenotypes.

**Fig. 2.**
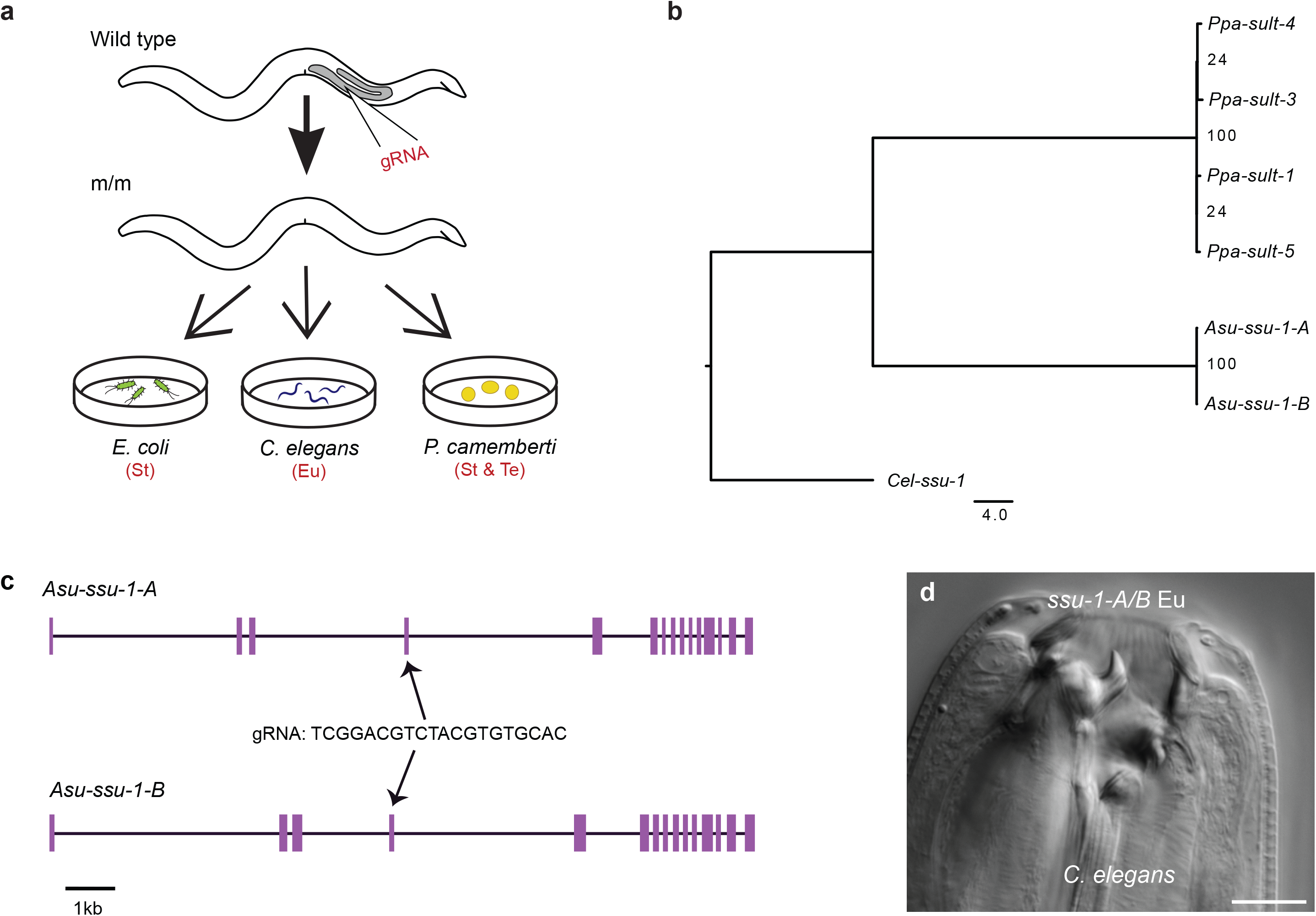
The sulfotransferase genes have a conserved role in Eu regulation. a) Wild type worms were injected with a gRNA targeting both genes via CRISPR. Homozygous mutants (m/m) were selected for and grown on 3 diets: 1) *E. coli* bacteria that induces St, 2) *C. elegans* that induces Eu and 3) *P. camemberti* that induces either St or the novel Te morph, all in wild type worms. b) The duplicate sulfotransferases *Asu-ssu-1-A* and *Asu-ssu-1-B* are homologous to four *P. pacificus* sulfotransferases, based on amino acid sequences, including *Ppa-sult-1* that has been shown to act as a switch in Eu regulation. c) The predicted gene structure of *Asu-ssu-1-A* and *Asu-ssu-1-B* shows high similarity as expected after WGD. A gRNA was designed that was able to target the predicted exon 4 in both genes. d) The *Asu-ssu-1-A/B* double mutant knock-out became Eu instead of St on *E. coli*, which only produces the St morph in wild type worms.

### Two sulfotransferase genes act as a switch in the regulation of the St morph

In *P. pacificus*, a cytosolic sulfotransferase-encoding gene, *sult-1*, homologous to *C. elegans ssu-1*, plays a key role as a switch gene. Knock-outs in *sult-1* prevent the narrow-mouthed St morph from forming, with mutants instead displaying the wide-mouthed Eu morph (Fig. 1b) (Namdeo *et al*. 2018; Bui *et al*. 2018). This mutant phenotype is thus the opposite of that seen in mutants of the sulfatase-encoding gene *eud-1* with mutants in *P. pacificus* being St under all conditions (Ragsdale *et al*. 2013). We previously showed the *A. sudhausi* sulfatase homologs of *P. pacificus eud-1*, termed *Asu-sul-2-A and Asu-sul-2-B*, together act as switch genes, with knock-out lines becoming St (Wighard et al. 2024) (Fig. 1g). We therefore wanted to determine if homologous *A. sudhausi* sulfotransferases also retain their switch gene function. Two homologs of *sult-1* had already been identified in *A. sudhausi, ssu-1-A* and *ssu-1-B* (Biddle and Ragsdale 2020) (Fig. 2b), which show high sequence similarity (Fig. 2c) and are presumably duplicates resulting from the recent *A. sudhausi*-specific WGD (Wighard et al. 2022).

We targeted *A. sudhausi ssu-1-A* and *ssu-1-B* and obtained single frameshift knock-out mutants using CRISPR; however, they displayed no mouth-form phenotype differences. This result was expected as previous studies already indicate most *A. sudhausi* duplicates resulting from WGD display redundancy (Wighard et al. 2022). We then crossed the single mutants together to obtain a double mutant and grew them on the three diets, as previously described. Strikingly, the *ssu-1-A/B* double mutant displayed an Eu phenotype on the St-inducing *E. coli* condition (Fig. 2d). This is similar to the phenotype of *P. pacificus sult-1* mutants which also remain Eu (Namdeo *et al*. 2018; Bui *et al*. 2018). On *P. camemberti, ssu-1-A/B* appeared able to become Te (Fig. S1), but did not form the St morph whatsoever. Thus, the *ssu-1-A* and *ssu-1-B* genes appear to regulate the St phenotype. This finding indicates that there is conservation of the role of the sulfotransferase genes in St regulation between both *A. sudhausi* and *P. pacificus*.

Note that the *ssu-1-A/B* double mutant line was noticeably unhealthy, with low survival and difficulty to culture. Specifically, *ssu-1-A/B* mutants had lower brood size (Fig. S2) and smaller body size (Fig. S3), with longer generation times and cannibalistic behaviour also observed. This meant very few worms survived bleaching, which is the standard way to obtain eggs on the three diets and prevent contamination. The *ssu-1-A/B* line was therefore not used for further downstream mouth-form analysis. The sickly phenotype of the mutant line could be due to the negative effects of preventing sulfotransferase activity. Cytosolic sulfotransferases have a major role in detoxification as they promote excretion of xenobiotics (Kurogi *et al*. 2024). Therefore, in the absence of these sulfotransferases, compounds can accumulate in cells and cause toxicity that produce such detrimental effects. In contrast, *P. pacificus* has multiple cytosolic sulfotransferase, meaning *sult-1* mutants are likely saved from having this detrimental phenotype due to high redundancy (Fig. 2b) (Igreja and Sommer 2022).

### *nag* mutants display an intermediate mouth phenotype

Next, we examined homologs of the *P. pacificus* supergene locus that has a crucial role in mouth-form regulation (Sieriebriennikov *et al*. 2018) (Fig. 3a). This locus contains *eud-1* as well as two recently duplicated genes, *nag-1* and n*ag-2* that encode N-acetylglucosaminidases. Knock-outs in both *nag* genes in *P. pacificus* lead to constitutive expression of Eu, with the St morph no longer formed. These duplicates are in tandem orientation and located close to *eud-1*. Interestingly, the supergene locus is not present in *A. sudhausi*, as it formed after the ancestors of *P. pacificus* and *A. sudhausi* diverged (Biddle and Ragsdale 2020) (Fig. 3b). The *nag* duplication also only occurred in *P. pacificus* and its close relatives after the divergence from *Allodiplogaster*. It is thus compelling to examine whether the role of the ancestral *nag* gene in mouth-form regulation preceded or followed formation of the supergene locus. There are three possibilities: 1) the ancestral *nag* was already involved in mouth-form regulation before becoming part of a supergene locus, 2) the ancestral *nag* played no role in mouth-form regulation prior to its duplication, meaning it evolved this function along with becoming part of a supergene, or 3) the ancestral *nag* was not involved in mouth-form regulation but independently co-evolved this function in both *A. sudhausi* and *P. pacificus*; although this latter possibility is less likely it cannot be ruled out.

**Fig. 3.**
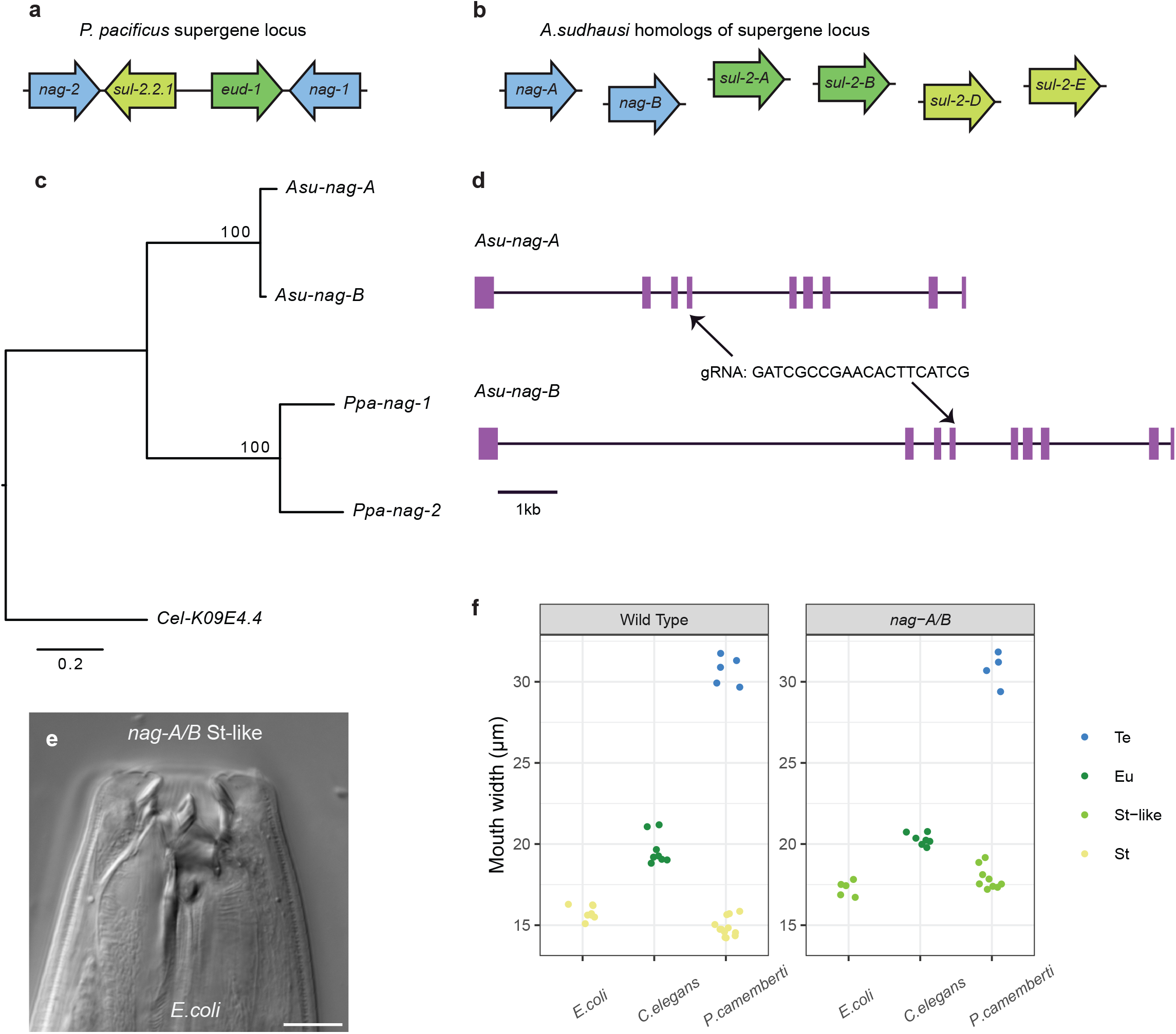
Knocking out *nag* genes produces an intermediate St phenotype. a) The *P. pacificus* supergene locus contains the *nag-1* and *nag-2* genes in tandem orientation surrounding sulfatase encoding genes, including *eud-1* which is involved in mouth-form regulation and its gene duplicate *sul-2*.*2*.*1*, which plays no role in mouth-form. b) *A. sudhausi* does not have a super gene locus. The homologs of those found in *P. pacificus* are found on separate contigs. c) The two duplicate *A. sudhausi nag-A* and *nag-B* genes are homologous to the *P. pacificus nag* genes that underwent a local gene duplication after *A. sudhausi* and *P. pacificus* diverged. d) The predicted gene structure is similar between *Asu-nag-A* and *Asu-nag-B*, with the same predicted number of exons. One gRNA was used to target exon 3 of both genes. e - f) On *E. coli*, the *A. sudhausi nag-A/B* double mutant knockout forms an intermediate ‘St-like’ phenotype whose mouth width is between wild type St and Eu, unlike the wild type which only becomes St on *E. coli*. The narrow morph from *E. coli* and occasionally *P. camemberti* diets is referred as either ‘St’ or ‘St-like’ on the x axis, depending on its respective mouth width.

We identified two *nag* homologs in *A. sudhausi*, which we termed *nag-A* and *nag-B* based on previous nomenclature (Fig. 3c). As the *Pristionchus* nag duplication took place after divergence from *Allodiplogaster*, the two *nag* genes in *A. sudhausi* would be WGD-derived duplicates of the single ancestral *nag* gene (Fig. 3d). Again, we found single mutants displayed a wild type phenotype across all diets, indicating redundancy in mouth-form function. Notably, the *nag-A/B* double mutant did not display the wild type St phenotype (Fig. 3e). Instead, *nag-A/B* displayed an intermediate ‘St-like’ phenotype, with the mouth width lying in-between that of wild type St and Eu morphs. Although the double mutant did not form the regular St phenotype on either *E. coli* or *P. camemberti*, it could still become Eu on *C. elegans* and Te on *P. camemberti* (Fig. 3f). This intermediate phenotype suggests *A. sudhausi nag-A* and *nag-B* genes together have quantitative effects. The phenotype in *A. sudhausi* thus differs from that in *P. pacificus*. In *P. pacificus*, the *nag* genes redundantly contribute to the switch, whereas in *A. sudhausi* they appear to be involved in the execution of the St morph and display a quantitative effect on mouth form. Importantly, the *nag* genes are involved in St regulation in both *A. sudhausi* and *P. pacificus*, suggesting *nag* genes have an ancestral role in formation of the St phenotype in Diplogastridae. Therefore, the involvement of the *nag* genes in mouth-form plasticity most likely preceded formation of the supergene locus.

### *nhr-40* mutants produce an intermediate phenotype of the novel Te morph

Comparisons between mouth-form genes in *A. sudhausi* and *P. pacificus* have thus far led to 1) conservation of function alone (*Ppa-sult-1, Asu-ssu-1-A/B*), 2) conservation of function and gain of novel function (*Ppa-eud-1, Asu-sul-2-A/B)*, and 3) partial conservation of function, with retention of St regulation in both species, but only acting as a switch in *P. pacificus* (*Ppa-nag-1/2, Asu-nag-A/B*). Next, we examined the nuclear hormone receptor-encoding gene *nhr-40*. This gene acts as a switch in *P. pacificus*, with loss of function *nhr-40* mutants preventing the Eu morph from being formed (Sieriebriennikov *et al*. 2020), similar to *Ppa-eud-1* mutants (Fig. 1b). We identified two WGD-derived homologous gene candidates in *A. sudhausi*, which we termed *Asu-nhr-40-A* and *Asu-nhr-40-B* (Fig. 4a). We again obtained frameshift knock-out mutant lines using CRISPR and examined their phenotypes on the three diets.

**Fig. 4.**
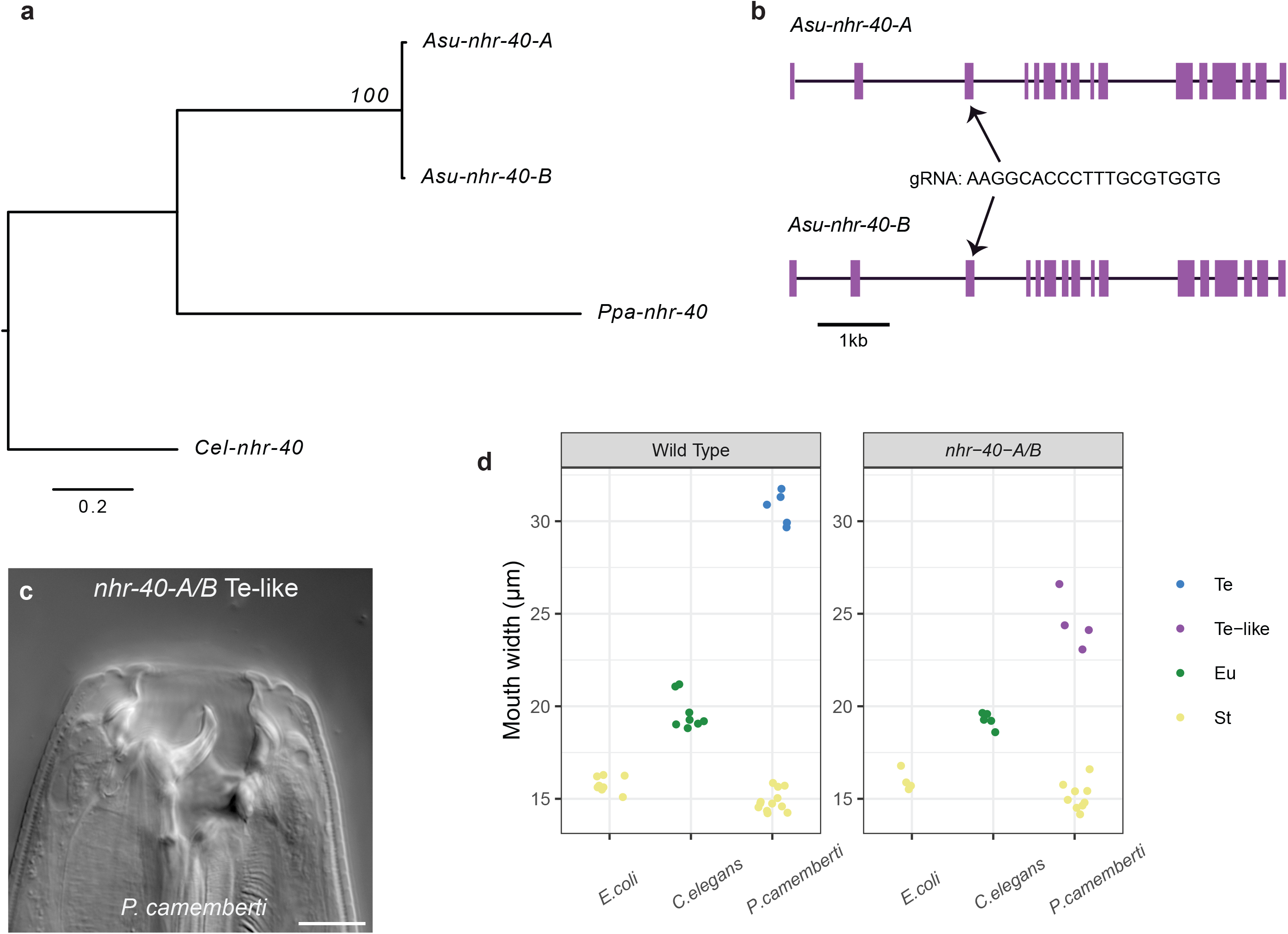
Knocking out *nhr-40* genes produces an intermediate Te phenotype. a) *Asu-nhr-40-A* and *Asu-nhr-40-B* were identified as the homologs of *Ppa-nhr-40*, with *nhr-40* as an outgroup. b) The gene structure is highly similar between the *A. sudhausi nhr-40* genes. A gRNA was designed that targeted exon 3 of both genes. c) The *nhr-40-A/B* double mutant knockout does not produce the regular Te phenotype on *P. camemberti*; instead, it forms a narrower ‘Te-like’ mouth that is otherwise similar in morphology. d) The nhr-40-A/B ‘Te-like’ is noticeable narrower than wild type Te and falls in-between the Eu and Te mouth width. The largest mouth-form found on *P. camemberti* was referred to either as ‘Te’ or ‘Te-like’ depending on mouth width.

Like the other WGD genes, *Asu-nhr-40-A* and *Asu-nhr-40-B* show high similarity in gene sequence and structure (Fig. 4b). The single knock-outs mutants again displayed a wild type phenotype, with gene redundancy occurring. However, the double mutant knock-out *nhr-40-A/B* displayed a clear yet unexpected mouth-form phenotype. The *nhr-40-A/B* knock-out line developed an intermediate ‘Te-like’ phenotype on the *P. camemberti* diet (Fig. 4c,d). That is, the mouth width lay in-between that of the Eu and Te phenotype, suggesting the genes have quantitative effects. In contrast, adults could still become St and Eu, as seen in wild type worms (Fig. 4d). Therefore, the *nhr-40-A* and *nhr-40-B* genes appear to be involved in regulating the novel Te phenotype. This is in stark contrast to *P. pacificus* where *nhr-40* acts as a switch gene in Eu regulation. Thus, the *nhr-40* genes in *A. sudhausi* firstly differ in that they are involved in regulation of a different mouth phenotype (Eu *vs*. Te) and, secondly, differ in that they do not act as a switch gene. Therefore, there is high divergence in the way *nhr-40* genes regulate mouth-form formation between *A. sudhausi* and *P. pacificus*.

The findings of the *A. sudhausi nhr-40-A/B* mutant can be compared to the *nag-A/B* results. There is a similarity in that, in both cases, the *A. sudhausi* homologs do not function as a switch and instead appear to have quantitative effects. However, there is a marked difference in the targets of the genes; the *nag* genes regulate the St phenotype in both *A. sudhausi* and *P. pacificus*, while the *nhr-40* genes regulate Eu in *P. pacificus* and Te in *A. sudhausi*. This unexpected finding is beneficial in that it allows us to formulate hypotheses on the timing of a particular evolutionary novelty in *A. sudhausi*. We previously showed that both WGD and evolution of the novel Te morph occurred after *A. sudhausi* diverged from its closest known relatives (Wighard et al. 2024); however, the timing of the Te morph and WGD relative to each other could not then be determined. Here, we show the *A. sudhausi* WGD-derived *nhr-40* genes are involved in formation of the Te mouth-form. This suggests that the Te morph evolved before WGD, as it is unlikely the *nhr-40* genes would both independently evolve to regulate this new morph. Thus, the evolution of the third morph might have preceded WGD in the lineage leading to *A. sudhausi* (but see a more detailed discussion below).

### The *nag* and *nhr-40* genes function independently of one another

Thus far, all homologous gene candidates vary in their degree of conservation of mouth-form function over large evolutionary distances (Table 1). This suggests there are multiple ways genes can functionally diverge. However, while the functional roles of genes have been compared between *A. sudhausi* and *P. pacificus*, the regulatory network and epistatic interactions of these genes had not been shown in *A. sudhausi*. We therefore generated combined *A. sudhausi* knock-out mutants. We first examined the relationship of *nhr-40/A/B* and *nag-A/B* genes to each other as their respective double mutant knockouts resulted in intermediate mouth-forms (Fig. 3e,f; Fig. 4c,d). These mutants display contrasting phenotypes because they regulate different mouth-form morphs, making them ideal to study their genetic relationship to one another. For instance, it could be epistatic, where knocking out one gene pair masks the expression of the other, or alternatively, they could function independently of one another. We injected the gRNA targeting both *nhr-40-A* and *nhr-40-B* into the *nag-A/B* double mutant and obtained a *nhr-40/A/B, nag-A/B* quadruple mutant knock-out (Fig. 5a). Strikingly, this quadruple mutant displayed the phenotypes of both the *nag-A/B* and the *nhr-40-A/B* double mutant lines (Fig. 5b). Specifically, on St-inducing *E. coli*, the quadruple mutant formed the intermediate wider St phenotype that was also seen in *nag-A/B* on the same diet. Similarly, on Te-inducing *P. camemberti*, the quadruple mutant formed the intermediate narrow Te mouth that was seen in *nhr-40-A/B* on the *P. camemberti* diet. No wild type St or Te mouth-forms were observed whatsoever in the quadruple mutant. Therefore, knocking out all four genes did not affect the phenotype of the other double mutant. We thus conclude that the *nhr-40-A/B* and *nag-A/B* genes function independently of each other, in parallel pathways of the mouth-form gene regulatory network (GRN).

**Table 1:** Homologous genes and their manner of conservation between *P. pacificus* and *A. sudhausi*. S: Switch genes; Q: Quantitative effects.

| Homologous mouth-form genes in diplogastrids |  | Role in mouth-form plasticity |  | Manner in which gene is conserved or diverged |  |
| --- | --- | --- | --- | --- | --- |
| *P. pacificus* gene name(s) | *A. sudhausi* gene names | Role in *P.pacificus* | Role in *A.sudhausi* | Conservation | Divergence |
| *eud-1* | *sul-2-A* & *sul-2-B* | Regulate Eu (S) | Regulate Eu (S)<br>Regulate Te (S) | Switch gene in Eu regulation | Novel function as switch gene in Te morph |
| *sult-1/seud-1* | *ssu-1-A* & *ssu-1-B* | Regulate St (S) | Regulate St (S) | Switch gene in St regulation | / |
| *nag-1* & *nag-2* | *nag-A* & *nag-B* | Regulate St (S) | Regulate St (Q) | Regulation of St morph | Switch genes in *P. pacificus* ; additive effects in *A. sudhausi* |
| *nhr-40* | *nhr-40-A* & *nhr-40-B* | Regulate Eu (S) | Regulate Te (Q) | / | Novel function in Te regulation |

**Fig. 5.**
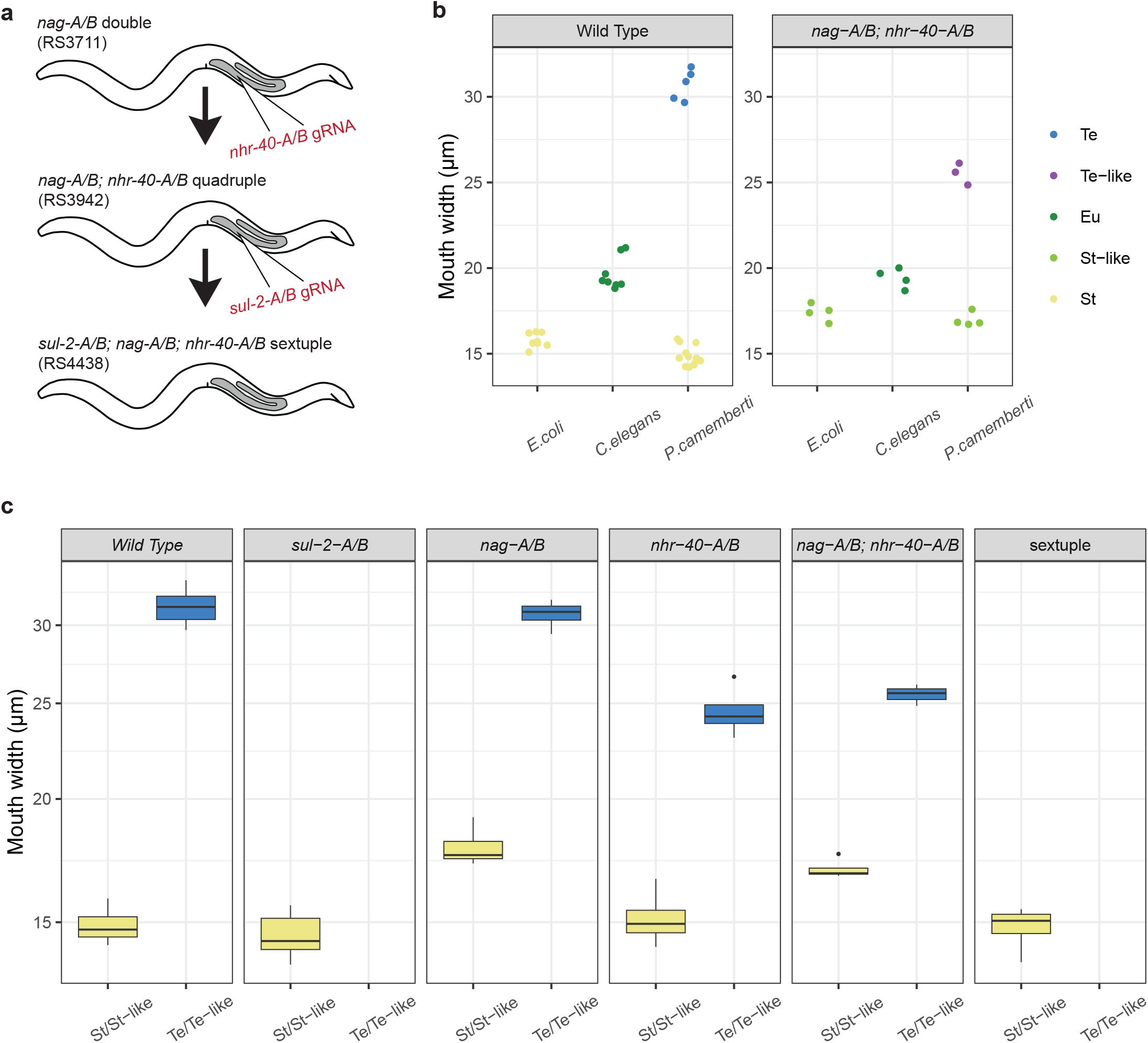
Combined mutations of mouth-form genes show epistasis in *A. sudhausi*. a) The gRNA that targets both *nhr-40-A* and *nhr-40-B* was injected into the *A. sudhausi nag-A/B* double mutant line to obtain the *nhr-40/A/B; nag-A/B* quadruple mutant knock-out. The gRNA that targets *sul-2-A* and *sul-2-B* was then injected into the quadruple mutant to obtain the *sul-2-A/B; nag-A/B; nag-A/B* sextuple knockout mutant. b) The *nhr-40/A/B, nag-A/B* quadruple mutant knock-out showed both the ‘St-like’ phenotype of *nag-A/B* on St-inducing diets as well as the ‘Te-like’ phenotype of *nhr-40-A/B*, implying these gene pairs function independently of one another. c) The mouth width of all examined mutant lines grown on a *P. camemberti* diet is shown. This diet can produce either St or Te adults in wild type. The mutants are shown next to one-another for comparison. The sextuple mutant remained St, indicating *sul-2-A/B* is epistatic to both the *nag* and *nhr-40* genes in *A. sudhausi*.

### The sulfatase genes act downstream of *nhr-40* and *nag* genes

Lastly, we examined the relationship of the sulfatase switch genes to *nhr-40* and *nag* genes. *Asu-sul-2-A/B* mutants are known to be constitutively St under all conditions (Wighard et al. 2024). We injected the *sul-2-A/B* gRNA into the *nhr-40/A/B; nag-A/B* quadruple mutant line to obtain a *sul-2-A/B*; *nhr-40/A/B; nag-A/B* sextuple mutant (Fig. 5a). Then, we grew the sextuple mutant line on St-inducing *E. coli*. On this diet, *nag-A/B* forms an intermediate wider St mouth, while *sul-2-A/B* forms the regular St mouth-form. Notably, the sextuple mutant line displayed the St phenotype (Fig. S4; Table S1), indicating the sulfatase genes are epistatic to the *nag* genes. Interestingly, this is also the case in *P. pacificus*, showing conservation of the gene regulatory pathway between both species.

We then grew the sextuple mutant on Te-inducing *P. camemberti*. The sulfatase double mutant is consistently St on all conditions; while the *nhr-40* mutant can form the narrower Te-like morph. Strikingly, we found the *A. sudhausi* sextuple mutant did not become Te at all (Fig. 5c; Fig S3; Table S1), indicating *sul-2-A/-B* acts downstream of *nhr-40-A/B*. Indeed, the sextuple mutant remains St under all conditions. Therefore, the sulfatase genes act downstream to the other mouth-form genes studied. This order in the GRN contrasts with the hypothesis in *P. pacificus* that *nhr-40-A/B* is epistatic to *sul-2-A/B*. While knockouts of the homologs in *P. pacificus* have the same phenotype, epistatic interactions are harder to interpret; although *Ppa-nhr-40 gain-of-function* alleles isolated from *eud-1* suppressor screens suggest both genes act in parallel to each other (Kieninger et al. 2016). The findings in *A. sudhausi* thus provide insight into the gene regulatory GRN in diplogastrids and its epistatic interactions.

## Conclusion

In this work, we examined homologous genes (based on sequence conservation) between two highly diverged nematode species. Remarkably, all candidate genes retained a role in mouth-form regulation in both *A. sudhausi* and *P. pacificus*. However, the exact role of these genes was not always conserved and, instead, displayed distinct identities of functional diversification in both species (Table 1). These results indicate there are multiple ways genes can functionally diverge over time, suggesting the complex mouth-form GRN is subject to continued selection at the gene-specific level. Indeed, the differences in the means of conservation may be due to relaxed pressure that allowed different pathways or parts of the GRN to evolve distinctively. Nevertheless, selection for mouth-form plasticity must be strong as the same homologous genes are involved in mouth-form regulation despite the two species last sharing a common ancestor nearly 200 million years ago (Qing et al. 2023). This suggests plasticity is a key process for survival of diplogastrid worms and their continued evolution. Therefore, our work on the evolution of plasticity in two distantly related species supports the hypothesis of plasticity as a major facilitator of evolutionary novelty (West-Eberhard 2003).

While each of the homologous gene pairs in *A. sudhausi* showed different means of conservation, it is particularly interesting that only the genes encoding the sulfatases and sulfotransferases retain their function as switch genes across both species. This suggests that the sulfation pathway is indeed crucial for regulating mouth-form plasticity within the Diplogastridae family. Nonetheless, further work is necessary to elucidate this pathway and better understand how and why sulfation processes are involved in nematode mouth-form plasticity (Igreja and Sommer 2022). Specifically, ongoing work tries to identify the target molecules of sulfation processes, *i*.*e*. hormones or other signaling molecules.

Interestingly, in examining conservation of the *nhr-40* homologs, we can provide first hypotheses regarding the previously held mystery of the relative timing of major evolutionary phenomena in *A. sudhausi*.

Originally, we identified the evolution of both the novel Te morph and the incidence of WGD (Wighard et al. 2024); however, the chronology of these events could not be determined due to a sufficient lack of closely-related strains. Here, we suggest that the Te morph evolved before WGD occurred, as the *Asu-nhr-40-A/B* mutant did not form the wild type Te phenotype (Fig. 4c,d). It is highly unlikely these gene would independently gain the same novel function in parallel. Therefore, since both *nhr-40* genes display the same function, it suggests WGD should have taken place after gain of the Te morph. Alternatively, the critical event for the evolution of the Te morph required the evolution of new target gene(s) after WGD. For instance, a potential scenario is that a new binding site for NHR-40 evolved, meaning there is instead a novel set of target genes that produced this functional shift. Thus, the results obtained in this study allow the formulation of two alternative hypotheses that can be further addressed in future studies. Note that WGD was estimated to have taken place only 1.3 to 3.3 million years ago (mya) (Wighard et al. 2022), meaning that the evolution of the Te morph happened in the time scale of a few million years.

In summary, we show that these mouth-form genes have ancestral functions in feeding structure plasticity throughout the Diplogastridae, but their specific roles have diverged in different lineages. Although *A. sudhausi* may be seen as the more ‘ancestral’ species due to its phylogenetic position, both species are identical in evolutionary age, meaning the species most similar to the common ancestor cannot be determined with certainty unless these genes are investigated in a broader range of diplogastrid. Taken together, our study highlights the dynamic nature of gene function divergence over time, adding to the body of work on sequence-level evolutionary changes. The plasticity network in diplogastrid nematodes presents an ideal model for studying gene divergence through evolutionary time, and we hope this study provides valuable insights for future research.

## Supporting information

Data S1: Sequences

Data S2: Mouth width measurements

## Data availability

Strains are available upon request. The authors affirm that all data necessary for confirming the conclusions of the article are present within the article, figures, and tables.

## Acknowledgements

We would like to thank all members of the Sommer Lab for the support, particularly Drs. Adrian Streit and Catia Igreja.

## Materials and Methods

### Nematode maintenance and inbreeding

The *A. sudhausi* inbred wild type strain SB413B/RS6132B (Wighard et al. 2022) and RS398 (*sul-2-A/B* double mutant knock-out) (Wighard et al. 2024) line was used. Further strains were generated via CRISPR knock-out engineering (described in detail further below): RS3711 (*nag-A/B* double mutant knock-out), RS3723 (*nhr-40-A/B* double mutant knock-out), RS3942 (*nag-A/B; nhr-40-A/B* quadruple mutant knock-out), RS4438 (*sul-2-A/B; nag-A/B; nhr-40-A/B* sextuple mutant knock-out). *C. elegans* (N2) was used as an Eu-inducing diet. All strains are frozen and kept at the Max Planck Institute for Biology. All nematodes were maintained on nematode growth medium (NGM) agar plates with *E. coli* OP50 bacteria.

### Nematode diets

Freshly starved NGM *E. coli* plates containing many eggs were bleached (Hope 1999) onto separate NGM containing three different food sources: 1) *E*.*coli* OP50, 2) *C. elegans* N2 and 3) *P. camemberti*, as previously described (Wighard et al. 2024). For *C. elegans*, plates with many larvae were washed with M9 and passed through a 20 µm nylon net filter (Merck) onto unseeded 6 cm NGM plates, so that only larvae passed through. *P. camemberti* was obtained from the Leipniz Institute DSMZ-German Collection of Microorganisms and Cell Cultures GmbH (https://www.dsmz.de) and maintained weekly on NGM plates. All assays were otherwise were maintained at 20°C. For all experiments, only young hermaphrodites were selected (containing less than five eggs) for mouth-form identification and measuring.

### Phylogenetic trees

The species tree was based on previous data (Susoy et al. 2015). For the gene trees, *P. pacificus* homologs were identified using sequence annotations from pristionchus.org (El Paco V3 2020). The corresponding protein sequences were blasted on wormbase.org (version WS284) to get *C. elegans* homologs. The predicted *A. sudhausi* protein sequences were identified on pristionchus.org using the previously published gene models (Wighard et al. 2022). The exact gene nomenclature and their corresponding annotations can be found in Table S2. The phylogenetic tree was generated using RAxML version 8.2.12 (raxmlHPC -f a -m PROTGAMMAAUTO -p 12345 -x 12345 -N 100), with maximum likelihood boostrap values included (raxmlHPC -f b -m PROTGAMMAILG) (Stamatakis 2014). The phylogeny was then displayed using FigTree software (v.1.4.4) (tree.bio.ed.ac.uk/software/figtree). The amino acid sequences of all predicted proteins can be found in Data S1.

### Microscopy images and mouth measurements

Young adult hermaphrodites were fixed onto a solution with 5% Noble Agar and 0.3% NaN3, with M9 buffer for resuspension. They were imaged at 100x magnification using the Differential Interference Contrast (DIC) setting. Only worms fixed in the correct orientation (on their lateral side) were picked. Z-stacks were taken of the head region and stored as raw data .czi files. Fiji/ImageJ (Schindelin et al. 2012) was used to measure the width of the mouth (stoma), using the base of the cheilostom and the start of the gymnostom the markers. All worms taken from the same assay plate were counted as one biological replicate. At least three biological replicates were used for each diet and corresponding mouth-form per strain. The exact measurements for each replicate can be found in Data S2.

### Mouth-form phenotyping

Worms were counted and phenotyped under a Zeiss Stemi 508 light microscope when they reached adulthood. The mouth-form of adult worms was then determined from all three assays.

### Brood size and worm size measurements

To compare wild type and *ssu-1-A/B* double mutants, both brood size and worm length were measured. For brood size, young virgin hermaphrodites were moved from well-fed *E. coli* plates to NGM plates containing 30 µl of *E. coli*. They were transferred to fresh plates ever two days for a total of eight days. The number of live offspring that each hermaphrodite produced was counted. All plates were stored at 20°C. For worm sizing, young adult hermaphrodites from *E. coli* were moved to NGM plates without bacteria. Bright-field images were taken at 20X magnification using the Axio Zoom V16 microscope. Analysis was done on Fiji/ImageJ using the Wormsizer plugin for ImageJ/Fiji (Moore et al. 2013) to detect worm length.

### CRISPR injections to obtain mutants

CRISPR knockouts were generated as previously described (Wighard et al. 2022). CRISPR RNAs (crRNAs) and trans-activating crispr RNA (tracrRNA) (Cat. No. 1072534) were synthesized by Integrated DNA Technologies (IDT), while the Cas9 endonuclease (Cat. No. 1081058) was purchased from IDT. The CRISPR/Cas9 complex was prepared by mixing 0.5 mg/ml Cas9 nuclease, 0.1 mg/ml tracrRNA, and 0.056 mg/ml crRNA in the TE buffer followed by a 10-min incubation at 37° C. Microinjections were performed in late-stage J4 hermaphrodites using an Eppendorf microinjection system. A single guide sequence was designed to target the two gene duplicates for each homologous gene pair of interest. Conserved regions were therefore targeted. Specific primers were designed for each individual gene in *A. sudhausi* (Table S3). Polymerase chain reactions (PCRs) were run using these primers on the F1 of injected worms. Heterozygotes were identified via Sanger sequencing (GENEWIZ Germany GmbH) and homozygous mutants were generated by selfing heterozygotes. Frameshift mutants were obtained for all strains. For *A. sudhausi ssu-1* homologs, single mutants were obtained from the first CRISPR, which were then crossed to get the *ssu-1-A/B* double mutant. Single mutant *nag-A* and *nag-B* were also gained from one CRISPR injection, and then a second injection was performed to get the *nag-A-B* double mutant. The *nhr-40-A/B* double mutant line was obtained after a single injection, as were independent single *nhr-40-A & -B* mutants. The *nhr-40-A/B* gRNA was injected into the *nag-A/B* double mutant line and a quadruple mutant was found after one injection. The *sul-2-A/B* gRNA (Wighard et al. 2024) was injected into the *nag-A/B; nag-A/B* quadruple mutant line, which resulted in the *sul-2-A/B; nag-A/B; nag-A/B* sextuple mutant after a single injection. Exact strain designations are shown above.

