## Supplementary Materials for "Functional divergence of conserved developmental plasticity genes between two distantly related nematodes"

**Affiliations:**

**This file includes:**

Figures S1 to S4

Tables S1 to S3

**Other Supplementary Materials include the following:**

Data S1 to S2

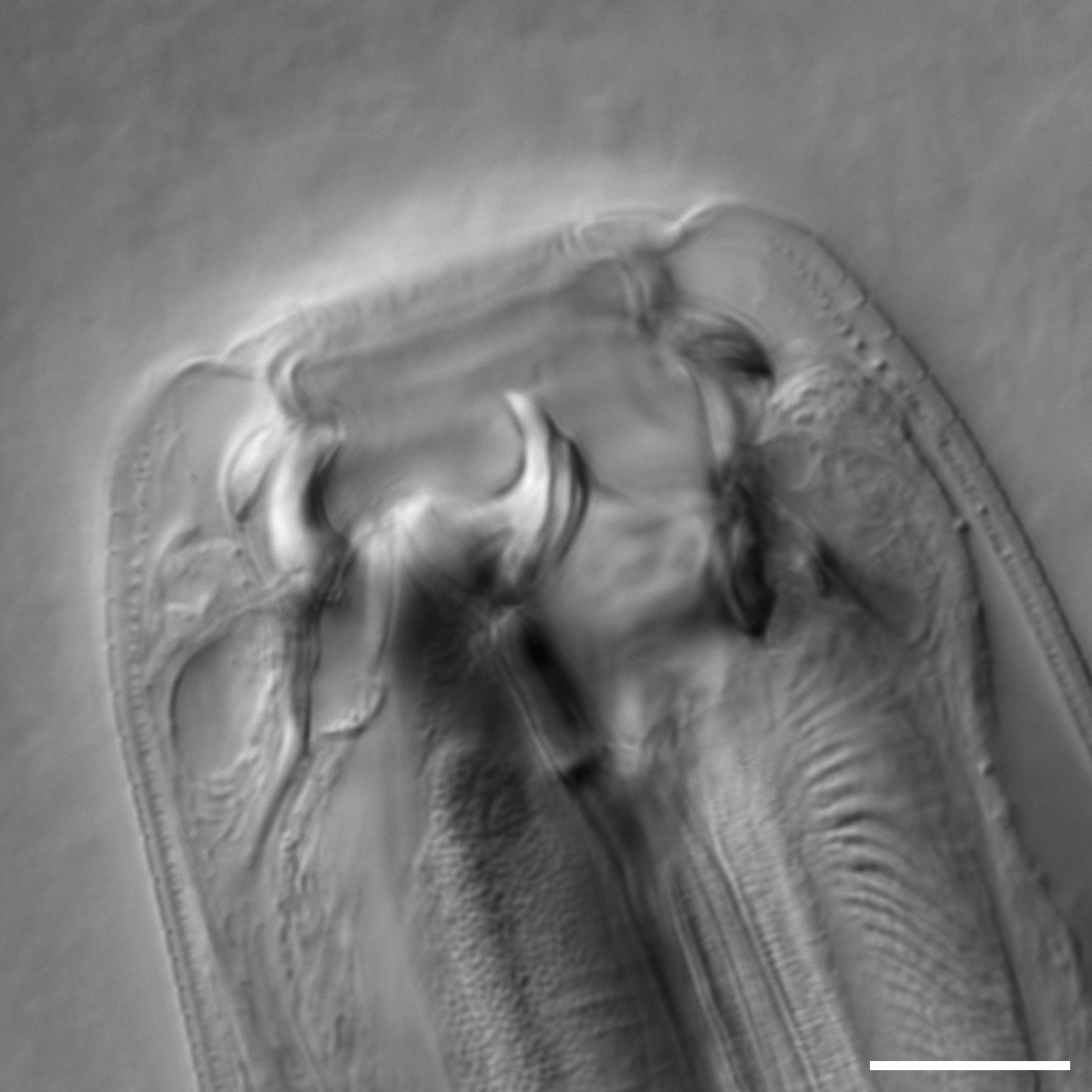

**Figure S1**: The *ssu-1-A/B* double mutant knockout can become Te on *P. camemberti* as seen in this DIC image. Scale bar: 10 μm.

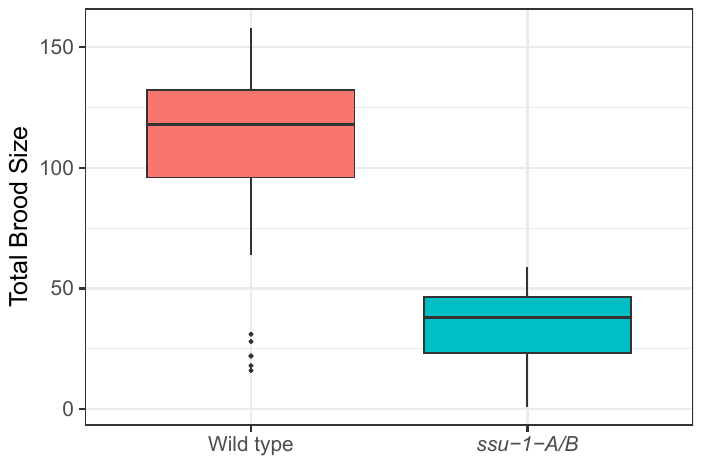

**Figure S2**: The *ssu-1-A/B* double mutant has a drastically lower brood size compared to wild type worms.

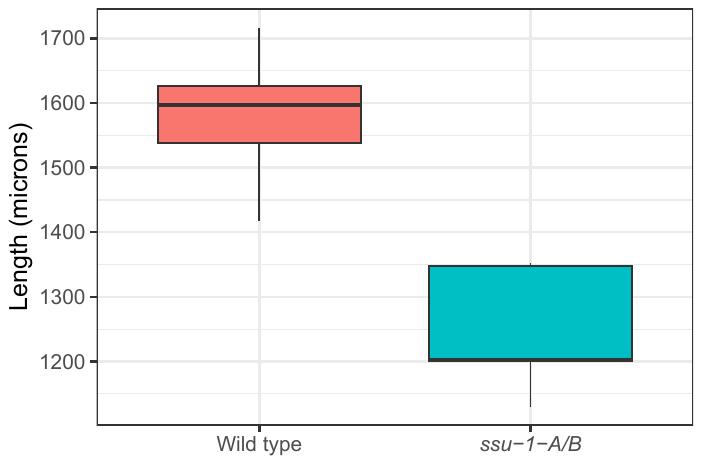

**Figure S3**: The *ssu-1-A/B* double mutant has a notably smaller body length compared to wild type worms.

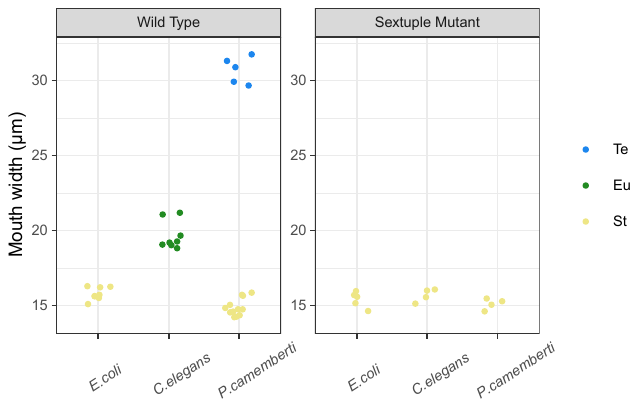

**Figure S4**: The *sul-2-A/B*; *nhr-40/A/B; nag-A/B* sextuple mutant remains St on all three diets and display a narrow mouth width, in contrast to wild type worms which become Eu on *C. elegans* and can become Te on *P. camemberti.*

**Table S1**: Phenotyping of the *sul-2-A/B*; *nhr-40/A/B; nag-A/B* sextuple mutant adult mouth-form on *C. elegans and P. camemberti* diets show they are consistently St.

| Diet | Replicate | No. St | No. Eu | No. Te | Total no. worms |
| --- | --- | --- | --- | --- | --- |
| *C. elegans* | 1 | 6 | 0 | 0 | 6 |
| *C. elegans* | 2 | 4 | 0 | 0 | 4 |
| *C. elegans* | 3 | 11 | 0 | 0 | 11 |
| *C. elegans* | 4 | 12 | 0 | 0 | 12 |
| *P. camemberti* | 1 | 6 | 0 | 0 | 6 |
| *P. camemberti* | 2 | 8 | 0 | 0 | 8 |
| *P. camemberti* | 3 | 33 | 0 | 0 | 33 |
| *P. camemberti* | 4 | 25 | 0 | 0 | 33 |
| *P. camemberti* | 5 | 55 | 0 | 0 | 55 |
| *P. camemberti* | 6 | 82 | 0 | 0 | 82 |
| *P. camemberti* | 7 | 66 | 0 | 0 | 66 |

**Table S2:** The gene names and respective annotations of the amino acid sequences, which can be found on pristionchus.org for *A. sudhausi* (Wighard *et al.,* 2022) and *P. pacificus* (El Paco annotation v3, 2020).

| Species | Gene name | Annotation |
| --- | --- | --- |
| *A. sudhausi* | *Asu-nag-A* | ALDISUDHAUS000014281 |
| *A. sudhausi* | *Asu-nag-B* | ALDISUDHAUS000017088 |
| *A. sudhausi* | *Asu_nhr-40-A* | ALDISUDHAUS000009481 |
| *A. sudhausi* | *Asu_nhr-40-B* | ALDISUDHAUS000005589 |
| *A. sudhausi* | *Asu-ssu-1-A* | ALDISUDHAUS000003619 |
| *A. sudhausi* | *Asu-ssu-1-B* | ALDISUDHAUS000003344 |
| *P. pacificus* | *Ppa-nag-1* | PPA06134 |
| *P. pacificus* | *Ppa-nag-2* | PPA34489 |
| *P. pacificus* | *Ppa-nhr-40* | ppa_stranded_DN28158_c0_g3_i3 |
| *P. pacificus* | *Ppa-sult-1/seud-1* | PPA12547 |
| *P. pacificus* | *Ppa-sult-3* | PPA06620 |
| *P. pacificus* | *Ppa-sult-4* | PPA22156 |
| *P. pacificus* | *Ppa-sult-5* | PPA41942 |

**Table S3:** The primer sequences to amplify the specific gene duplicates and identify mutants after CRISPR injections.

| Primer name | Primer sequence (5' - 3') | Forward/Reverse | Target |
| --- | --- | --- | --- |
| sul-2-A_F | AGAATGTAGCCAGGCAAGC | Forward | *Asu-sul-2-A* |
| sul-2-A_R | CTCAGTCGACATGGAAAAGC | Reverse | *Asu-sul-2-A* |
| sul-2-B_F | ATTGCAGGATACGGCGACC | Forward | *Asu-sul-2-B* |
| sul-2-B_R | CTAATCTCGTCTATGCCGACG | Reverse | *Asu-sul-2-B* |
| ssu-1-A_F | TCTTCTTGCGACCAATGCGG | Forward | *Asu-ssu-1-A* |
| ssu-1-A_R | TAGAGCAGTCTGGACAAAGC | Reverse | *Asu-ssu-1-A* |
| ssu-1-B_F | TGATTGCGCGAAACGGAGAC | Forward | *Asu-ssu-1-B* |
| ssu-1-B_R | ATTAGAAGGATTGGCCGTGC | Reverse | *Asu-ssu-1-B* |
| nag-A_F | ATGGACTCGGTTACTTCACC | Forward | *Asu-nag-A* |
| nag-A_R | GACATTTGCTGGCTTCGTGC | Reverse | *Asu-nag-A* |
| nag-B_F | CAGTTCCACCAAACCATCG | Forward | *Asu-nag-B* |
| nag-B_R | TTGCTCCACCTCAATTTACG | Reverse | *Asu-nag-B* |
| nhr-40-A_F | CCTTAAATTAGGCTTTGAGC | Forward | *Asu-nhr-40-A* |
| nhr-40-A_R | CACTCTCCATTACCATACACT | Reverse | *Asu-nhr-40-A* |
| nhr-40-B_F | CTTGCGTAACTCCTTCTAATC | Forward | *Asu-nhr-40-B* |
| nhr-40-B_R | AGTATTGAGGTGAAGCTGGC | Reverse | *Asu-nhr-40-B* |
